# Population differences in aggression are shaped by cyclone-induced selection

**DOI:** 10.1101/612838

**Authors:** Alexander G. Little, David N. Fisher, Thomas W. Schoener, Jonathan N. Pruitt

## Abstract

Surprisingly little is known about the evolutionary impacts of rare but extreme black swan events, like tropical cyclones. By intercepting three cyclones in fall 2018, we evaluated cyclone-induced selection on collective behavior in a group-living spider. We further examined whether historic frequencies of cyclone landfalls are correlated with geographic variation in group behavior. Cyclones consistently selected for more aggressive spider societies. Furthermore, sites where cyclones have historically been more common also harbor more aggressive groups. Thus, two corroborative lines of evidence convey that that cyclone-induced selection can drive the evolution of colony behavior, and suggest that extreme black swan events can shape within-species diversity and local adaptation.

**One Sentence Summary:** Tropical cyclones drive the evolution of more aggressive spider societies.

## Main Text

Extreme events, such as natural disasters, are destructive yet influential ecological forces (*1*). However, observing and recording the effect of these extreme “black swan” events (*2, 3*) is logistically difficult, and so rigorous examination of their roles in ecological and evolutionary processes is lacking. Tropical cyclones, for instance, represent some of the most impressive forces of nature on planet Earth. High wind speeds, erosion, precipitation, and storm surge from cyclones can transform familiar habitats into desolate landscapes for their constituents. While the effects of cyclones are well quantified in terms of human wellbeing and economic costs (*4*), their ecological and evolutionary consequences for non-human populations are less clear.

Habitat alterations caused by cyclones promise to radically alter the selective pressures acting on many organisms. Raging winds, for instance, can demolish trees, defoliate entire canopies and scatter coarse woody debris across the forest floor (*5, 6*). Correlative studies on post-cyclone landscapes convey that they alter the behavior (e.g. (*7*)), nutrition (e.g. (*8*)), abundance (e.g. (*9*)), population demographics (e.g. (*10*)), and metapopulation connectivity (e.g. (*11*)) of many organisms. Given the diversity and scales of these effects, it stands to reason that the evolutionary and ecological dynamics of cyclone-prone environments will be starkly divergent from environments that do not experience these extreme events (*12-14*). While intuitively appealing, there is little direct evidence that this is the case. This is because cyclone ecology is largely characterized by opportunistic, *post-hoc* studies that lack spatiotemporal replication and reference (control) sites in their design. Cyclones occur with seasonal regulatory, but landfall positioning, timing, and intensity can be difficult to forecast on a storm-by-storm basis. This unpredictability makes the rigorous examination of cyclone ecology logistically complicated, but not impossible. As the environmental impacts of cyclones only promise to grow as sea levels rise (*15*), the pressure to understand the environmental impacts of these storms has never been more pressing.

Multi-female colonies of the group-living spider (*Anelosimus studiosus*) occur along the Gulf and Atlantic coasts of the United States and Mexico. This places them in the path of cyclones formed in the Atlantic basin from May to November. *A. studiosus* produces cobwebs containing one to a few hundred females on the tips of branches overhanging bodies of water – most often streams and rivers (*16*). This species is characterized by a behavioural dimorphism, where individuals exhibit either a heritable docile or aggressive phenotype (*16*). Natural colonies of *A. studiosus* contain mixtures of docile and aggressive females, and the relative representation of each phenotype additively determines the collective aggressiveness of a colony (*17, 18*). Colony aggressiveness determines the speed and number of attackers that respond to prey (*17, 19*), prey sharing efficiency and wastage (*19, 20*), tendency to cannibalize males and eggs (*16, 21*), and susceptibility to infiltration by predatory and parasitic foreign spiders (*22*). Like individual-level aggressiveness (*16*), colony aggressiveness in *A. studiosus* is transmitted down colony generations (*22*), from parent to daughter colony (*23*), and is a major determinant of spiders’ survival and fecundity in habitat-and site-specific manners (*16, 24*).

Here we use *A. studiosus* as a model to determine the intensity of cyclone-induced selection on colony behavior and evaluate whether the frequency of these rare but extreme black swan events can explain regional differences in colony behavior. A rigorous understanding of the evolutionary impacts of tropical cyclones has evaded the literature to date both because of the obvious logistical and methodological challenges of contending with giant storms, and because most of the available literature tends not to consider their evolutionary implications (but see (*25*)). Here we sampled 240 colonies of *A. studiosus* in cyclone-disturbed and paired control reference sites before and after each of three tropical cyclones: Subtropical Storm Alberto (N_sites_= 1 storm, 1 reference), Hurricane Florence (N_sites_= 2 storm, 1 reference), and Hurricane Michael (N_sites_= 3 storm, 1 reference). This was accomplished by traveling to sites within the projected trajectory of each storm and assaying the behavior and demographics of naturally-occurring colonies in the storm’s path at storm-strike sites, paired with a reference site that resided well outside the projected path of the storm but within the same general region. We returned to each site within 48 hours to confirm which colonies persisted through the storm. Following cyclones, we recorded the number of egg cases produced by each colony (October, 2018) and the number of spiderlings that survived until winter (December, 2018). Spiderlings disperse the following spring (*26*). Because aggressiveness is favored under low resource conditions in many spiders (*27-29*), and because flying insects (i.e,. prey) often increase in abundance following tropical cyclones (*30*), we hypothesized that cyclones would reduce selection for aggressiveness in *A. studiosus* and increase colonies’ reproductive output. If, however, cyclones are expensive in terms of added web repair costs or reduced prey availability, then we might predict cyclones to select for more aggressive societies.

We also sampled 211 colonies of *A. studiosus* at sites distributed across the American range of this species (Fig. 1a) in May-June 2018, prior to hurricane season, to test for associations between a history of cyclone strikes and regional variation in colony behavior. For all colonies we recorded (1) the height above water, (2) an estimate of web volume (L × W × H), (3) the diameter of the immediate supporting branch, (4) the number of attackers that responded to a vibratory cue in the web over two minutes (colony aggressiveness), and (5) the number of females in the colony (Full Methods: SI 1).

**Fig 1.**
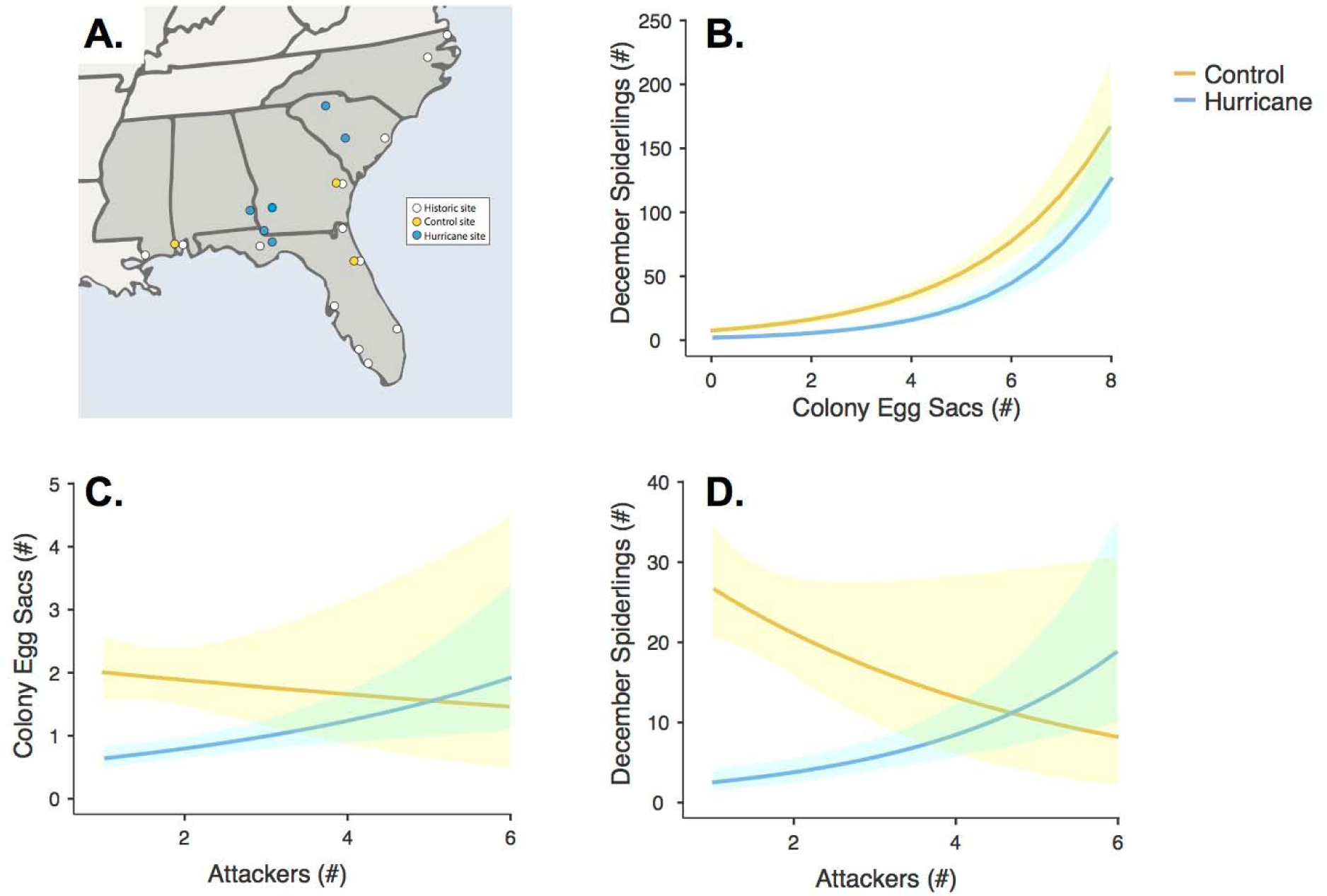
The effects of cyclones on colony fecundity. Effects of control (yellow) and cyclone-disturbed (blue) sites (A) on Decemeber spiderlings as a function of colony egg sac numbers (B), and the influence of number of attackers on colony egg sacs (C) and December spiderlings (D). Highlighted areas represent 95% CI.

At odds with our primary expectations, we found that cyclones reduced the number of egg cases produced by colonies (β=0.326; 95% CIs=0.146, 0.500; df=1; p<0.001) as well as the number of spiderlings that survived until the winter (β=0.622; 95% CIs=0.352, 0.900; df=1; p<0.001), relative to control sites (Fig. S1a,b). Cyclones further reduced the number of spiderlings that survived until winter per egg case produced (β=−0.0654; 95% CIs=−0.109, −0.0217; df=1; p<0.001; Fig. 1b), implying that cyclones reduced the survival of spiderlings long after the cyclone had dissipated. These negative impacts resonate with the few studies on disparate terrestrial species demonstrating that juveniles are particularly susceptible to cyclone strikes (*5, 31, 32*).

Cyclones selected for more aggressive colony phenotypes. Following cyclones, colonies with more aggressive foraging responses produced more egg cases (“cyclone” x “number of attackers” interaction: β=−0.1417; 95% CIs=−0.2768, −0.0192; df=1; p<0.031; Fig. 1c) and had more spiderlings survive into early winter (“cyclone” x “number of attackers” interaction: β=−0.3193; 95% CIs=−0.4836, −0.173; df=1; p<0.001; Fig. 1d), whereas the opposite trend emerged in control sites.

The importance of cyclone-induced selection on colony aggressiveness is further bolstered by the retrospective comparison across 13 populations. We found that regional variation in colony aggressiveness was correlated with the local history of cyclone disturbance events. The proportion of spiders that respond to prey was positively correlated with the number of cyclone strikes (category 1-5) in the county over the past 100 years (β= 0.0226; 95% CIs=0.0068, 0.0384; df=1; p=0.005; Fig. 2). This association was not driven by spatial autocorrelation (see SI 2, Fig S4). Thus, not only do cyclones appear to have a consistent pattern of selection on colony aggressiveness across multiple storms, but the importance of these storms is apparently strong enough to drive regional variation in a behavioral trait that is both heritable and functionally significant.

**Fig 2.**
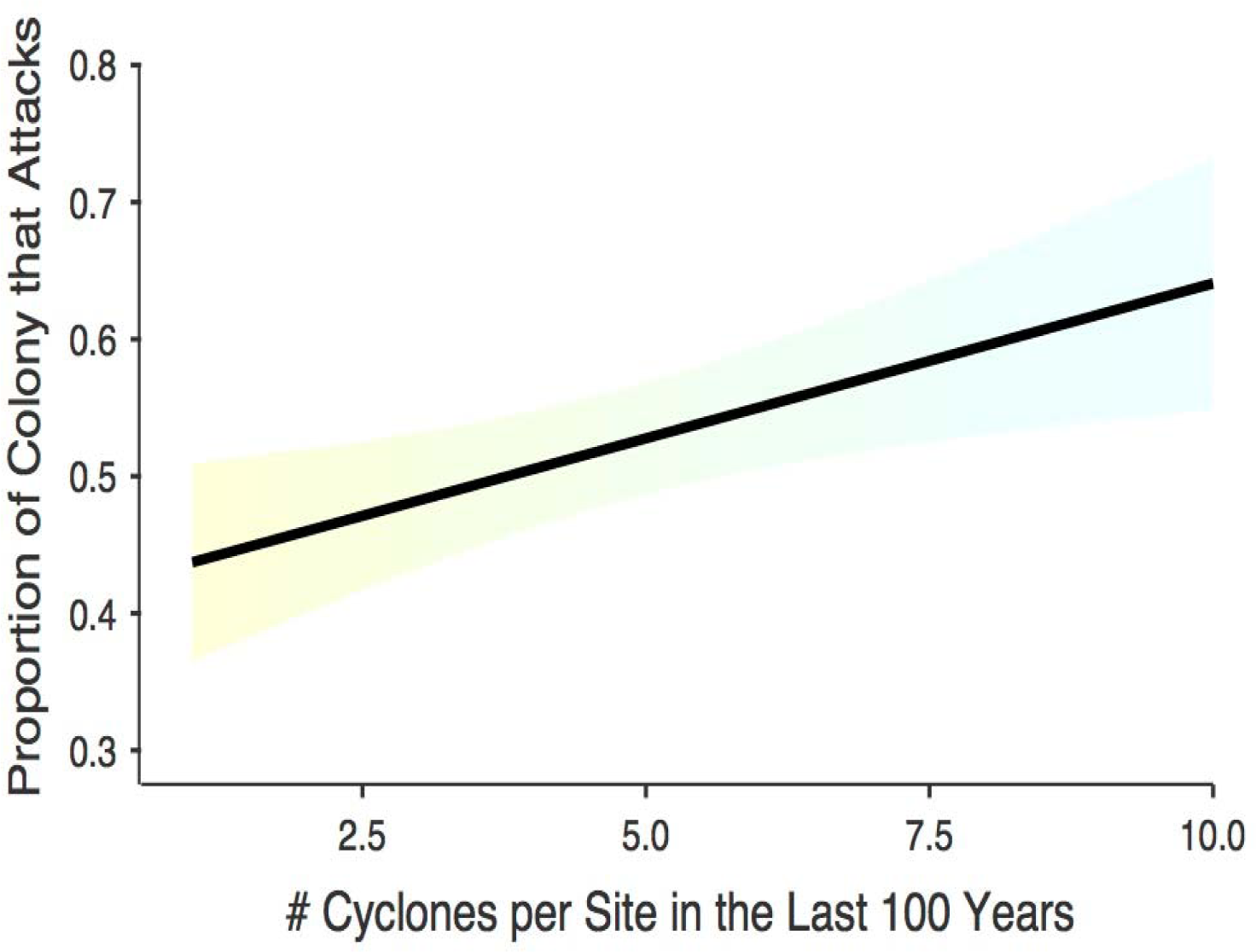
The effect of local cyclone frequencies on colony aggression. The relationship between the number of cyclones per site in the last 100 years and the proportion of a colony that attacks. Highlighted area reflects 95% CI, with yellow to blue gradation representing low to high frequencies of cyclones.

We do not presently understand the mechanism(s) explaining why aggressive colonies outperform docile ones following cyclone strikes. We reason that post-cyclone environments could drive down prey availability, which enhances selection on aggressiveness in web-building spiders (*27-29*). Post-cyclone environments may also increase the motility and aggressiveness of other species of spider, which might be more likely to invade *A. studiosus* colonies following cyclones. These intruders too are known to select for more aggressive colonies in *A. studiosus* (*18, 22*). Finally, cyclones may reduce the longevity of *A. studiosus* mothers engaged in parental care, which imperils the young in this species (*33, 34*). If this is the case, then it is possible that aggressive offspring may be less reliant on the prey and defense of their mothers, and therefore, suffer less mortality.

Our findings show that cyclones select for more aggressive colony phenotypes, and that this selection appears significant enough to drive regional variation in behavior. This is despite the fact the cyclone strikes represent irregular events, occurring only every few years, even in particularly cyclone-prone regions. While fecundity and offspring survival are reduced by cyclone disturbance in *A. studiosus*, colonies with proportionally more attackers have higher reproductive outputs, a trend that is not true of control sites. This trend is consistent across multiple storms that varied in both size, duration, and intensity. This too conveys that these effects are not merely idiosyncratic, and instead, are robust responses that hold across storms and at sites occupying a spread of more than 5° latitude. In aggregate, these data provide the most definitive evidence to date for cyclone-induced selection driving the evolution of an important functional trait and convey that black swan events contribute to within-species diversity and local adaptation.

## Author contributions

Conceptualization: A.G.L and J.N.P. Supervision: J.N.P. and T.W.S. Investigation and Methodology: J.N.P. and A.G.L. Formal Analysis and Visualization: A.G.L. and D.N.F. Writing -original draft: A.G.L. Writing -review and editing: A.G.L, D.N.F., T.W.S. and J.N.P.

## Data and materials availability

All raw data is available in a supplementary data file.

## Supplementary Materials

Materials and Methods

Figures S1-S4

Data S1

References (*35-37*)

